# N^6^-methyladenosine regulates Influenza A virus mRNA stability yet is rarely found on genomic RNA

**DOI:** 10.64898/2026.09.01.748482

**Authors:** Hsiu-Yi Wu, Wan-Ju Tung, Ramreishang Wungmaiwo, Pei-Yi (Alma) Su, Hung-Wei Hsu, Ankit Gupta, Brian N Papas, Marcos Morgan, Wen-Chun Liu, I-Hsuan Wang, Kevin Tsai

**Affiliations:** Institute of Biomedical Sciences (IBMS), Academia Sinica, Taipei, Taiwan; Taiwan International Graduate Program in Molecular Medicine (TIGP), National Yang-Ming Chiao-Tung University and Academia Sinica, Taipei, Taiwan; Reproductive and Developmental Biology Laboratory, National Institute of Environmental Health Sciences, National Institutes of Health, Durham, NC, USA; Integrative Bioinformatics, Biostatistics and Computational Biology Branch, National Institute of Environmental Health Sciences, National Institutes of Health, Durham, NC, USA; Infectious Disease Core Facility, Biomedical Translation Research Center, Academia Sinica, Taipei, Taiwan; Department of Microbiology and Immunology, College of Medicine, Kaohsiung Medical University, Kaohsiung, Taiwan

## Abstract

Previous studies have found widespread N^6^-methyladenosine (m^6^A methylation) on all forms of Influenza A virus (IAV) RNA, with m^6^A found critical for viral replication, pathogenicity as well as viral RNA packaging. Here we applied the latest quantitative technologies to revisit the methylation landscape on the anti-sense genomic RNA of IAV. Unexpectedly, upon Ultra-Performance Liquid Chromatography-Tandem Mass Spectrometry (UPLC-MS/MS) analysis of IAV virion -extracted genomic RNA, we detected very little m^6^A regardless of production from human cells or chicken eggs. Concordantly, Nanopore direct RNA sequencing also detected an overall low occurrence and stoichiometry (generally <5%) of m^6^A across all viral genomic RNA segments, compared with abundant m^6^A sites on viral mRNAs at ∼20-30% m^6^A. Cross validation with glyoxal- and nitrite-mediated deamination of unmethylated adenosines (GLORI) confirmed multiple m^6^A sites on viral mRNA yet very few m^6^A on the genomic RNA. This paucity of m^6^A on genomic RNA makes it unlikely that m^6^A contributes to viral RNA packaging. Knockdown or pharmacological inhibition of the m^6^A methyltransferase METTL3 as well as the reader protein YTHDF2 both reduced viral mRNA levels and infectious viral particle production, with YTHDF2 promoting viral mRNA stability. Thus, the presence of m^6^A on IAV transcripts is indeed proviral, yet it is the mRNAs instead of genomic RNAs that are methylated at functionally relevant levels. Lastly, we provide proof of concept that a METTL3 small molecule inhibitor can be antiviral, and propose that m^6^A-targeted antivirals would mainly impact the intracellular gene expression phase of IAV replication.

**Importance:** Influenza A virus (IAV) is a contagious pathogen that cause seasonal epidemics, and with a large reservoir among birds, has historically switched hosts on multiple occasions and contributed to several pandemics. The rapid evolution of influenza viruses necessitates the development of antivirals that work across virus strains, and one approach is to target host mechanisms that are needed for the replication of a wide range of viruses. The adenosine methylation m^6^A, is present on the RNA of a wide range of viruses including IAV, where it mostly supports viral replication. However, it remains unclear how abundant is m^6^A on IAV RNA. Here, we utilize the latest RNA modification detection methods and found that IAV mRNAs are m^6^A methylated, with little m^6^A on the genomic RNA. We provide evidence that m^6^A on IAV mRNAs may prevent degradation of IAV mRNAs, and provide proof of concept that an m^6^A modification inhibitor can be antiviral against IAV.

## Introduction

RNA within cells are subject to small chemical modifications, such as methylations and acetylations, that regulate the fate and function of the modified RNA (1). Also known as epitranscriptomic RNA modifications, the most abundant of these modifications on mammalian mRNAs is the methylation of the N6 position of the Adenosine base, termed N^6^-methyladenosine (m^6^A), with m^6^A reported to regulate the splicing, nuclear export and stability of the methylated mRNA (2–4). As obligate parasites, the RNA transcripts of viruses are also subject to the same methylations as host mRNAs, with m^6^A found to regulate the RNA of a wide variety of viruses (5). In particular, we and others have found m^6^A to enhance the RNA stability of Human Immunodeficiency Virus type 1 (HIV-1) and Hepatitis B virus (HBV) (6–8), support the nuclear export of the mRNA of Simian Virus 40 (SV40) (9), and prevent the viral RNA of Respiratory Syncytial virus (RSV) and Human Metapneumovirus (hMPV) from activating cellular interferon responses (10), illustrating an evolutionary convergence of diverse viruses utilizing m^6^A for enhancing viral replication and fitness.

Influenza A virus (IAV) was among the first three viruses whose RNA transcripts were found to be m^6^A methylated. IAV belongs to the Orthomyxoviridae family, with viral particles containing eight segments of a negative-sense single stranded RNA genome, each genomic RNA segment coated with nucleoprotein (NP) and a trimeric viral RNA polymerase complex forming viral ribonuclear protein complexes (vRNP) (11). Upon infection, the eight vRNPs enter the host cell nucleus and serve as the template for production of sense-strand viral mRNAs as well as replication into progeny anti-sense viral genomic RNA, all through the utilization of viral RNA dependent RNA polymerase complexes. In a pioneering report in 1976, Krug et al. analyzed the RNA methylation content under conditions where ∼90% of the intracellular poly(A)+ RNA in infected cells are IAV mRNAs, and found that 55-60% of the methyl (CH_3_) content on sense-strand IAV mRNAs are in m^6^A (12). The presence of m^6^A on IAV mRNA was later confirmed, with varying levels of m^6^A found on different viral mRNA transcripts (13). However, the function of such methylations remained unclear while it was also unknown if the negative-strand genomic RNAs were methylated or not. More recently, technological advances in high throughput sequencing and the development of m^6^A RNA-immunoprecipitation (meRIP) enabled us to map m^6^A on both sense and antisense IAV transcripts (14). The addition of m^6^A onto IAV transcripts was dependent on the cellular methyltransferase METTL3, with m^6^A enhancing the gene expression, replication and virulence of IAV, likely through the recruitment of the host m^6^A reader protein YTHDF2 which is generally thought to act as an adapter between m^6^A+ RNA and RNA-effector enzymes (3). However, it remains unclear how m^6^A enhances IAV replication. One recent report concurred that m^6^A acts as a pro-viral factor and reported that m^6^A methylation of the IAV genomic RNA was required for genome packaging into RNPs since mutational removal of meRIP-identified m^6^A sites from the viral negative strand genomic RNA greatly reduced RNP packaging (15).

While meRIP has greatly enhanced our understanding of how m^6^A regulates mRNAs and viral RNAs alike, limitations of meRIP-derived m^6^A-mapping strategies have gradually became apparent, with the methylated site within the pulled down ∼100nt RNA fragment unclear, false positive sites pulled down due to non-specific antibody binding and/or insufficient washes, and unclear m^6^A stoichiometry (16). While our initial study sought to minimize these limitations through photoactivated crosslinking of the m^6^A antibody to viral RNA incorporated with the uridine analogue 4-thiouridine (photo-crosslinking-assisted m^6^A-seq, PA-m^6^A-seq) (14, 17), the resulting m^6^A map lacked stoichiometry information, thus leaving it unclear if each site was methylated on every copy of the viral transcript or was limited to a few rare copies of that transcript. Recent technological advances have allowed us to better tackle the stoichiometry issue, with UPLC-MS/MS allowing us to accurately quantify the relative amount of methylated versus unmethylated adenosine bases (18), albeit with no information on the location of the methylated adenosines. Alternatively, Nanopore direct RNA sequencing (DRS) allows us to directly identify the sequence and modifications of an RNA molecule by threading the RNA through a protein pore and recording the disruption of ion currents passing through that pore, and matching that electrical signal to a pre-trained machine learning model to identify the sequence of A, U, C, Gs as well as m^6^As (19, 20). Furthermore, m^6^A can also be mapped at single nucleotide resolution through the selective deamination of unmethylated adenosines using a mixture of glyoxal and nitrite, leaving m^6^As intact. Subsequent reverse transcription of this deaminated RNA followed by high-throughput sequencing enables the identification and quantification of the chemically-untouched m^6^As, in a method termed GLORI (21).

Here, we exploit the latest methods including UPLC-MS/MS, Nanopore DRS, and GLORI to revisit the m^6^A epitranscriptomic landscape of IAV transcripts. Unexpectedly, UPLC-MS/MS analysis of viral particle-extracted genomic RNA suggested that m^6^A is present at levels >10x lower than that typically seen on cellular mRNAs, an observation that is consistent across human cell line or embryonated egg -produced virion-packaged RNA. While m^6^A-methylated negative-strand viral RNA could indeed be immunoprecipitated during prior meRIP-associated assays, Nanopore DRS and GLORI suggested that the m^6^A content on IAV genomic RNA is only present at vanishingly low stoichiometries across four tested H1N1 and H3N2 strains. While artificial production of fully m^6^A-methylated IAV genomic RNA enhanced RNA binding with viral NP protein by ∼25%, we found no evidence that a forced increase in m^6^A rates could increase virus production in cells. Rather, m^6^A was consistently found on the sense-strand viral mRNAs by RNA IP, Nanopore and GLORI, where we found that m^6^A to mainly support the stability of IAV mRNAs through the recruitment of the cellular m^6^A reader protein YTHDF2, and demonstrate that pharmacological inhibition of m^6^A addition is a viable antiviral strategy for IAV. Thus, we argue that m^6^A is a pro-viral host factor for IAV replication, and that this is mainly contributed through the m^6^A methylation of viral sense-strand mRNAs rather than the anti-sense genomic RNAs.

## Methods

### Cells

Madin Darby Canine Kidney (MDCK) epithelial cells, DF-1 chicken embryonic epithelial cells, A549 human lung adenocarcinoma cells and 293T human kidney epithelial cells were cultured in Dulbecco’s Modified Eagle’s Medium (DMEM) with 10% FBS and 1% Antibiotic-Antimycotic (Corning #30-004-CI). DF-1 cells were maintained at 39℃ while all other cell lines grown at 37℃. DF-1 cells were a kind gift from Hui-Wen (Winni) Chen (Department of Veterinary Medicine, National Taiwan University). A549 cells with a tet-inducible METTL3 expression vector and METTL3-knockout A549 cells were kind gifts from Alexander Price (The Wistar Institute) and Matthew Weitzman (Children’s Hospital of Philadelphia) (22).

### Viruses

Virus clones used include A/Puerto Rico/8/1934 (H1N1, abbreviated as PR8) either produced in MDCK cells from eight bidirectional pDZ plasmids (23, 24) (kindly provided by Peter Palese, Icahn School of Medicine at Mount Sinai) or produced in 9-10 days-old embryonated chicken eggs inoculated with stock PR8 virus (American Type Culture Collection, ATCC). Strains A/WSN/1933(H1N1), A/Udorn/307/1972 (H3N2), A/Aichi/2/1968 (H3N2) (abbreviated WSN, Udorn and Aichi) were all similarly produced in embryonated chicken eggs using stock virus from ATCC. All chicken egg-derived virus stocks produced at the Infectious Disease Core Facility, Biomedical Translation Research Center, National Biotechnology Research Park (ID Core, BioTReC, NBRP), Academia Sinica. Reference viral genomes used for sequencing alignments include: PR8: Genbank #EF190971-EF190978; WSN: Genbank #LC333182-LC333189; Udorn: Genbank #CY009636-CY009643; and Aichi: ATCC #VR547.

### IAV Infection

A549 cells were seeded to 40% confluency into 6-well or 150 mm plates. The next day, cells were washed with PBS and infected in infection media (PBS supplemented with 8.35M CaCl_2_, 10M MgCl_2_, 0.25% BSA, 1% Antibiotic-Antimycotic) for 1hr, using 0.01 MOI for virus amplification, 0.1 MOI for spreading infections, 0.5-1 MOI for single round infections and 5 MOI for virion RNA analysis. A549 cells were then washed with PBS and cultured in recovery media (DMEM with 0.15% FBS, 0.35% BSA and 1% Antibiotic-Antimycotic). Infected cells or medium were collected at 16-24 hours post infection (hpi) for single round infection or 1-2 days post infection (dpi) for spreading infections or virion purification.

### Virion Purification

Chicken egg -derived viruses were produced as mentioned above. Viruses grown in A549 cells (human) were harvested from the supernatant of thirty 15cm plates after 2 days of infection (5 MOI). Virus-containing media were centrifuged at 2200g for 20mins at 4°C to clean the cell debris. The virus media was spun at 10800g 30mins, then layered onto a 20% sucrose cushion in NTE buffer (100 mM NaCl, 10 mM Tris-Cl (pH 7.4), 1 mM EDTA) and pelleted by ultracentrifugation at 112600g for 2hrs at 4°C in a SW28 rotor (Beckman Coulter). The concentrated virus was fractionated over a 30–60% sucrose gradient at 112,600g for 3hrs at 4°C in a SW41 rotor (Beckman Coulter). Each fraction was collected and pelleted at 112,600g for 90mins at 4°C in a TLA110 rotor (Beckman Coulter), and resuspended in 0.5mls of NTE buffer. Each pelleted fraction verified for the presence of viral products by RT-qPCR and Western blot.

### UPLC–MS/MS Sample Preparation

DF-1 (chicken) or A549 (human) cellular mRNA were extracted with Trizol and poly(A) purified using the Poly(A)Purist MAG Kit (Invitrogen). The resulting mRNA and virion-extracted RNA were further depleted of ribosomal RNA using an NEBNext rRNA depletion Kit v2 (NEB), with rRNA depletion verified by microfluidic electrophoresis using a Fragment Analyzer (Agilent). These purified RNAs were digested to single nucleotides with the Nucleoside Digestion Mix (NEB #M0649S) for 2hrs at 37°C and cleaned using a Microcon-10KDa filter (Millipore #MRCPRT010).

### PA-m^6^A-seq

Done as previously described (14, 25). Briefly, A549 cells were infected at 5MOI with PR8, WSN, Udorn, or Aichi virus for 4hrs, then the virus-containing supernatants removed and cells overlayed with recovery media (see above for IAV infections, with 0.15% FBS omitted) supplemented with 100µM 4SU and incubated for another 20hrs, the infected cells finally harvested and the RNA extracted with Trizol. Poly(A)+ RNA were enriched from each sample with Sera-mag oligo(dT) beads (Cytiva), and 11-13µg of poly(A)+ RNA used for immunoprecipitation with 7.5µg of m^6^A antibody (Synaptic Systems #202 003), and the antibody-RNA complexes UV crosslinked. The pulled down and end-repaired m^6^A+ RNA fragments were used for sequencing library preparation using the NEBNext small RNA sequencing kit for Illumina (NEB #E7330), and sequenced on Illumina NextSeq 500 or 2000 sequencers. Resulting reads were clipped of adaptor sequences using fastx_clipper -Q33 -v -l 15, converted to fasta files and duplicate reads collapsed with fastx_collapser. The cleaned reads were pre-aligned to the human genome hg19 with bowtie -p 8 -v 1 -a -m 9 --best –strata --chunkmbs 30000, and the unaligned reads then aligned to viral genomes with bowtie -p 8 -m 8 -v 1 -a --best --strata --chunkmbs 30000. Viral alignment was done once with the --norc setting for sense strand alignment, and again with --nofw for anti-sense alignment. Reads with T>C or A>G mutations were selected for downstream analysis, realigned to the viral genome with bowtie and sorted and indexed with samtools for visualization in IGV.

### Nanopore Direct RNA-seq (DRS)

Viral RNA from A549 cells infected at 1MOI were harvested 2dpi, either from supernatant virions or from cell lysates of twelve 15cm plates, both extracted with Trizol. Nanopore DRS sequencing libraries were prepared from 1-2µg of total cell lysate RNA or RNA extracted from sucrose gradient -purified virions (see virus purification above), following Oxford Nanopore Technologies’ direct RNA sequencing library prep kit (SQK-RNA004) with minor modifications. Each RNA sample were separately ligated to standard DeePlexiCon barcoded RT adaptors (RTAs) (26) to tag the poly(A)+ mRNAs while also ligated to modified DeePlexiCon RTAs where the 10dT splint-ligation portion was replaced with an IAV-genomic RNA 3’end (U12) -specific sequence as previously designed (27). After the initial RTA ligation and clean up, mRNA-specific-RTA ligated libraries and genomic-RNA-specific-RTA ligated libraries were mixed at a 3:1 ratio and combined into one single RLA ligation reaction for motor protein addition. All samples were loaded onto a Nanopore RNA flow cell (FLO-MIN004RA) and run on a MinION MK1B sequencer.

Sequencing data was demultiplexed using Deeplexicon (26). The Deeplexicon model was retrained with libraries generated using the RNA004 library preparation kit (SQK-RNA004) and run on RNA flow cells (FLO-MIN004RA). Each demultiplexed dataset was basecalled using Dorado v0.8.1 with the rna004_130bps_sup@v5.1.0 model, with the argument --modified-bases m6A_DRACH. Sequences were then aligned to viral genomes using minimap2 with the arguments -t 28 -ax splice -uf -k14. Unmapped, reverse-aligned, non-primary and supplementary alignments were discarded from the aligned bam file with samtools view -F 2304. Positive strand (mRNA) aligning reads were selected with samtools view -F 20 while negative strand genomic RNA aligning reads selected with samtools view -f 0×10. m^6^A peaks were called using modkit pileup with --motif GAC 1 -- motif AAC 1, with the resulting methyl-bed files converted into bedgraph files of peaks showing % methylation using: awk ‘{if (($10/($10+$18))>0.05) {print $1 “\t” $2 “\t” $3 “\t” $11}}’, thus discarding all sites with excessive modification no-call events (N_nocall_ > 5%). Per-read m^6^A call statistics were extracted using modkit extract, also with the arguments --motif GAC 1 --motif AAC 1, modification calls with >0.95 probability scores were considered true m^6^A detection events.

### GLORI-seq

Performed following the recently improved GLORI 2.0 one pot reaction with minor changes (21). RNA was extracted from A549 cells infected with IAV at an MOI of 5 and harvested at 24hpi, and the poly(A)+ RNA isolated with Sera-mag oligo(dT) beads (Cytiva #38152103010150). The resulting RNA was fragmented in 10mM ZnCl_2_, 10mM Tris-HCl pH7.0 at 94°C for 45secs and subjected to glyoxal-mediated deamination. Following deamination and deprotection, the RNA (along with untreated controls) was purified, dephosphorated and re-phosphorated. Sequencing libraries were prepared using the NEBNext small RNA sequencing kit for Illumina (NEB #E7330), and sequenced on an Illumina NextSeq 2000 sequencer. Data was analyzed with GLORI-tools as described (28), with only m^6^A calls in GAC or AAC motifs retained.

### RNAi-mediated gene knockdown

A549 cells were transfected with siRNA (OriGene SR309781 for siYTHDF2 and SR324578 for siMETTL3) using Lipofectamine RNAiMAX transfection reagent (Invitrogen #13778150), with the three siRNAs in each gene target -specific kit pre-mixed in a 1:1:1 ratio for transfection. Cells were reseeded 24hrs later and subject to infection 48hrs post transfection.

### m^6^A ELISA

EpiQuik m^6^A RNA Methylation Quantification Kit (Epigentek, P9005) was used for m^6^A concentration measurement following the manufacturer’s guidelines.

### RT-qPCR

RNAs in cytosol were extracted using Trizol, and viral RNAs in media were obtained using Trizol LS and Direct-zol RNA MiniPrep Plus (Zymo, R2072). 1.5μg of RNA was treated with DNaseI, and reverse transcribed with SuperScript III (Invitrogen, 18080044) using RT primers specific for poly(A)+ mRNAs (CCAGATCGTTCGAGTCGTttttttttttttt) or antisense IAV genomic RNAs (CCAGATCGTTCGAGTCGTagcaaaagcaggg) as previously described (29, 30). SYBR Green Master Mix (Applied Biosystems) was utilized for qPCR on a QuantStudio 3 (Applied Biosystems). All primers used as listed in Table S1.

### Viral RNA decay assay

A549 cells were seeded and transfected with siRNA using Lipofectamine RNAiMAX (Invitrogen) at 50% confluency. At 24 hours post-transfection, cells were reseeded to 40% confluency. At 48 hours post-transfection, cells were infected with the virus at an MOI of 1. At 24 hours post-infection, the culture medium was replaced with new recovery medium containing Baloxavir acid (MedChemExpress). Cells were harvested at 0, 2, 4.5, and 7 hours post-treatment, lysed in Trizol, and subjected to qPCR analysis.

### Antibodies

METTL3 (Novus Biologicals H00056339 & Proteintech 15073-1-AP), YTHDF2 (Abcam ab220163), GAPDH (GeneTex GTX41577 & Proteintech 60004-1-Ig), PB1 (GeneTex GTX125923), PA (GeneTex GTX125932), HA (GeneTex GTX127357), M1 (GeneTex GTX125928), M2 (Genetex GTX125951), NS1 (Genetex GTX633685), FLAG tag (clone M2, Sigma F3165), anti-mouse (Sigma A9044) and anti-rabbit (Sigma A6154 & Cytiva NA934).

### Plaque assay

MDCK cells were seeded at 1×10⁶ cells/well in 6-well plates 24hrs prior infection. Cells were subject to infection with 10-fold serial dilutions of virus. 1hr later, cells were washed once with 1 mL of PBS and subsequently overlaid with 4 mL of DMEM supplemented with 0.15% FBS, 0.35% BSA, 0.1% trypsin and 1% agarose. 72hrs later, cells were fixed with 4% formaldehyde for >2hrs and subsequently stained with crystal violet. Virus titers were calculated as plaque-forming units (PFU) per milliliter.

### In vitro transcription (IVT) and RNA pull-down assay

The HA and NS genome segments of PR8 were cloned in the antisense direction into pGEM-3zf+ (Promega) after a T7 promoter, generating the plasmids pGEM-PR8-HA and pGEM-PR8-NS. pGEM-PR8-HA and pGEM-PR8-NS were linearized and used as templates for *in vitro* transcription (IVT) using the MEGAscript™ T7 Transcription Kit (Invitrogen, AM1334) with or without ATP substituted with m^6^ATP to generate RNAs containing or lacking m⁶A modification. The IVT RNAs were purified using the RNA Clean & Concentrator-5 Kit (Zymo, R1016).

HEK293T cells were transfected with plasmids encoding the PR8 NP protein (pK-FLAG-NP). At 48 h post-transfection, cells were harvested and washed three times with cold PBS, then lysed in NET buffer (50mM Tris pH 7.5, 150mM NaCl, 5mM EDTA, and 0.5% NP-40). Cell lysates were incubated with anti-FLAG antibody at 4°C for 1 h, followed by incubation with Protein G Mag Sepharose Xtra beads (Cytiva, 28967070) at 4°C overnight. The beads were washed three times with NET buffer and once with pre-warmed PBS (37°C). Subsequently, beads were incubated with IVT RNA containing or lacking m⁶A at 37°C for 30mins. After incubation, beads were washed once with lysis buffer (20mM Tris–HCl, pH7.5, 500mM LiCl, 1mM EDTA, 5mM DTT, and 0.5% LiDS), followed by sequential washes with wash buffer #1 (20mM Tris–HCl, pH7.5, 500mM LiCl, 1mM EDTA, 5mM DTT, and 0.1% LiDS), wash buffer #2 (20mM Tris–HCl, pH7.5, 500mM LiCl, 1mM EDTA, 5mM DTT, and 0.02% NP-40), and wash buffer #3 (20mM Tris–HCl, pH7.5, 200mM LiCl, 1mM EDTA, 5mM DTT, and 0.02% NP-40). Finally, RNA was extracted directly from the beads using TRIzol, and the amount of pulled-down RNA was quantified by qPCR.

### STM2457 treatment and infection of A549 cells

A549 cells were seeded in 12-well plates at ∼2×10⁵ cells/well and pretreated with STM2457 (Sigma-Aldrich #SML3360) at concentrations of 5, 10, 20, or 30 µM for 24 h prior to infection. Cells were then infected with 0.1 MOI of PR8-strain virus. The viral inoculum was removed 1hr later, cells washed once with PBS and overlayed with recovery medium containing trypsin (0.25%, 1µl per 1ml medium; Gibco, #15050-065) and STM2457 at the corresponding concentrations. Cells were incubated for an additional 23hrs at 37°C. At the indicated time point, cells were washed once with PBS and harvested in TRIzol reagent for RNA extraction or in Laemmli buffer for protein analysis by Western blotting.

### Cell Viability Assay

A549 cells were seeded in 96-well plates. After 24 hours, the culture medium was replaced with 150µl of medium containing various concentrations of STM2457. At 48 hours post-treatment, the culture medium was aspirated and replaced with 100μl of alamarBlue reagent (Invitrogen). Cells were incubated in the dark at 37°C for 1hr. Subsequently, the supernatants were transferred to black plates, and fluorescence was measured using a microplate reader (excitation: 560nm, emission: 590nm).

## Results

### IAV genomic RNA adenosines are hypo-methylated

As a zoonotic family of viruses that can infect multiple host species with varying fitness, we initially hypothesized that IAV viral RNA produced in birds and mammals may be differentially methylated. To test this, we purified virions of the H1N1 strain A/Puerto Rico/8/34 (PR8) from embryonated chicken eggs as well as infected human A549 lung epithelial cell supernatants, over sucrose cushions. The corresponding chicken and human intracellular poly(A)+ RNA was also purified from infected DF-1 chicken embryo fibroblasts and human A549 cells. Virion or cellular mRNA extracted from these chicken and human sources were subsequently digested to single nucleotides and the modified and unmodified nucleotide content was quantified by ultraperformance liquid chromatography tandem mass spectrometry (UPLC-MS/MS, Fig. 1A-B), with single nucleoside standards used for nine commonly seen RNA modifications as well as the four standard nucleosides. Contrary to our initial hypothesis, we only observed some differences in the 2’O-methylations (Cm, Gm, Am) between chicken and human -sourced virion RNA, with not much obvious cross-species differences in any of the other assayed modifications. However, we unexpectedly found the stoichiometry of m^6^A on virion RNAs at ∼0.05%, at least 10x lower than the generally 0.3-0.6% m^6^A/A among host cellular mRNAs (18, 29, 31).

**Fig 1.**
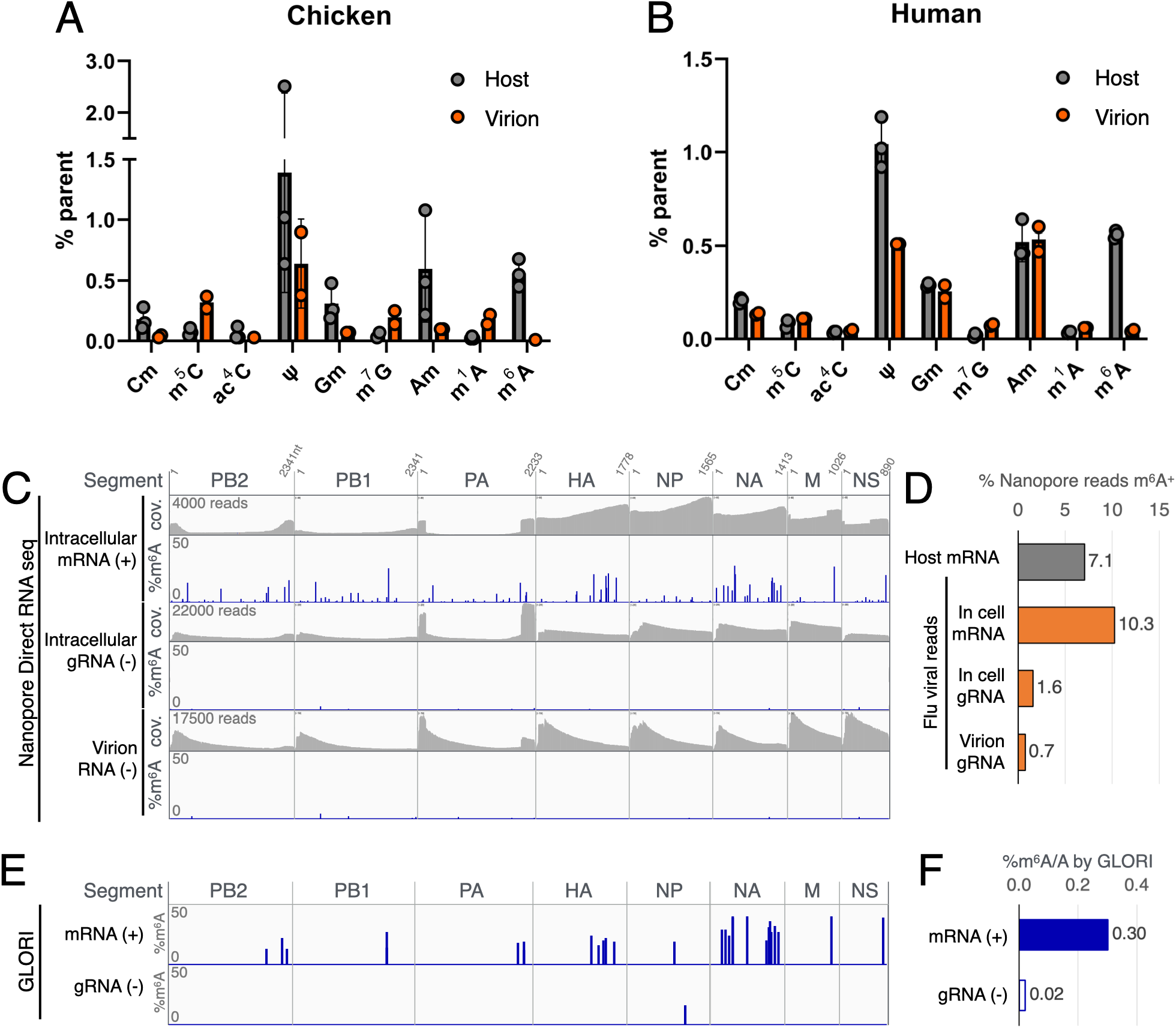
Influenza A viral genomic RNA is hypo-m^6^A-methylated. **(A-B)** Quantification of host mRNA and viral RNA modifications by UPLC-MS/MS, shown as % of parent nucleotide modified (e.g. %m^6^A/A for m^6^A). Measurements from infected chicken cellular total mRNA (DF-1 chicken embryo fibroblasts, “host”) and chicken egg -produced virion RNA shown in **(A)**, and infected human cell mRNA (A549 lung epithelial cells, “host”) and A549-produced virion RNA in **(B)**. **(C)** Viral RNA from infected A549 cells and the supernatant virions were subjected to Nanopore direct RNA seq (DRS), with the read coverage (cov.) and % of reads with detected m^6^A (%m^6^A) at each position shown. **(D)** Percentage of DRS reads in panel C with at least one m^6^A detected, comparing that of host mRNA, in cell viral mRNA, in cell antisense viral genomic RNA (gRNA) and virion-extracted gRNA. **(E)** Viral RNA from infected A549 cells were alternatively subject to GLORI detection of m^6^A, with the % m^6^A at each position shown separately for the sense strand mRNA (+) and antisense strand genomic RNA (gRNA(-)). **(F)** Calculated overall %m^6^A/A of IAV mRNA and gRNA from the GLORI data in panel E. All virus in this figure are of strain A/Puerto Rico/8/1934 (PR8).

As we have previously found m^6^A to be several fold enriched on the virion RNA of other viruses including HIV-1 and Hepatitis B virus, and m^6^A was previously found on IAV negative strand RNA using immunoprecipitation that was not very quantitative (14, 29, 32), we next seek to validate this unexpected observation using alternative methods that may inform us of the relative abundance of m^6^A on sense versus antisense IAV transcripts. For this purpose, we utilized Nanopore direct RNA sequencing (DRS), which identifies m^6^A on individual RNA molecules (33). Analysis of the m^6^A sites on viral RNA extracted from PR8-infected A549 cells as well as infected A549 supernatants using Nanopore DRS revealed that the sense strand mRNAs expressed from all eight segments can indeed be m^6^A methylated (Fig. 1C, lanes marked intracellular mRNA(+)), with most of these Nanopore-mapped m^6^A sites concurrent with previously immunoprecipitation-mapped m^6^A sites (Fig. S1). However, among the antisense viral RNA reads, regardless of those in cells or in supernatant virions, none of the m^6^A sites were present on more than 5% of the reads covering that specific site, with most peaks showing stoichiometries <2% (Fig. 1C bottom lanes marked (-)). When we quantified the number of DRS reads with at least one m^6^A site identified versus those with no methylation detected, we found the percent of m^6^A-positive reads among host and viral mRNAs comparable at 7.1 and 10.3% respectively, in stark contrast to that in the antisense genomic RNAs (in cell and virion) with the percent reads m^6^A+ at 0.7-1.6% (Fig. 1D). To ensure that the dearth of m^6^A detection on the negative strand viral RNAs were not due to Nanopore -specific sequencing artifacts, we alternatively utilized glyoxal- and nitrite-mediated deamination of unmethylated adenosines (GLORI) to map and quantify m^6^A on IAV RNA, where all but the m^6^A-methylated adenosines are deaminated into inosines and subsequently sequenced as guanosines (21). Upon performing GLORI on RNA extracted from PR8-infected A549 cells, we confirmed all the major peaks mapped on the IAV sense strand mRNA by Nanopore DRS, yet similarly noticed almost no m^6^A on antisense genomic RNAs (Figs. 1E & S1A), with the lone GLORI-detected antisense strand m^6^A curiously inconsistent with Nanopore DRS. As GLORI is quantitative, we also calculated the m^6^A/A ratio of sense and antisense IAV RNA, and found the viral sense strand RNA with 0.3% m^6^A/A rate close to that expected on mammalian mRNAs, while the low 0.02% m^6^A/A rate of antisense viral RNA is closer to the 0.01-0.05% detected in virion RNAs by UPLC-MS/MS (Figs. 1A-B & 1F).

### The lack of m6A on IAV genomic RNAs is consistent across multiple isolates

To ensure if our unexpected observation is not unique to the lab strain PR8, we expanded our Nanopore DRS analysis to three other influenza viral isolates in the lab, including the H1N1 strain A/WSN/1933 (WSN), and the H3N2 strains A/Udorn/307/1972 (Udorn) and A/Aichi/2/1968 (Aichi). Repeating our direct RNA sequencing on RNA extracted from A549 cells infected with PR8, WSN, Udorn, and Aichi, and separating the sense and antisense genome aligning reads (Fig. 2A-D), we again observed that the antisense genomic RNAs (gRNA(-)) of all strains tested contained m^6^A sites of much lower stoichiometry than that seen in the sense strand RNAs (mRNA(+)). Upon quantifying the number of reads with >1 m^6^A versus those devoid of m^6^A signals, we once again found across all four strains ∼8-14% of mRNA reads carry m^6^A signals, while only 0.4-0.5% of the antisense genomic RNAs (gRNA) are m^6^A+ (Fig. 2E-H). Thus, hypomethylation of IAV genomic RNAs is universal across multiple viral strains.

**Fig 2.**
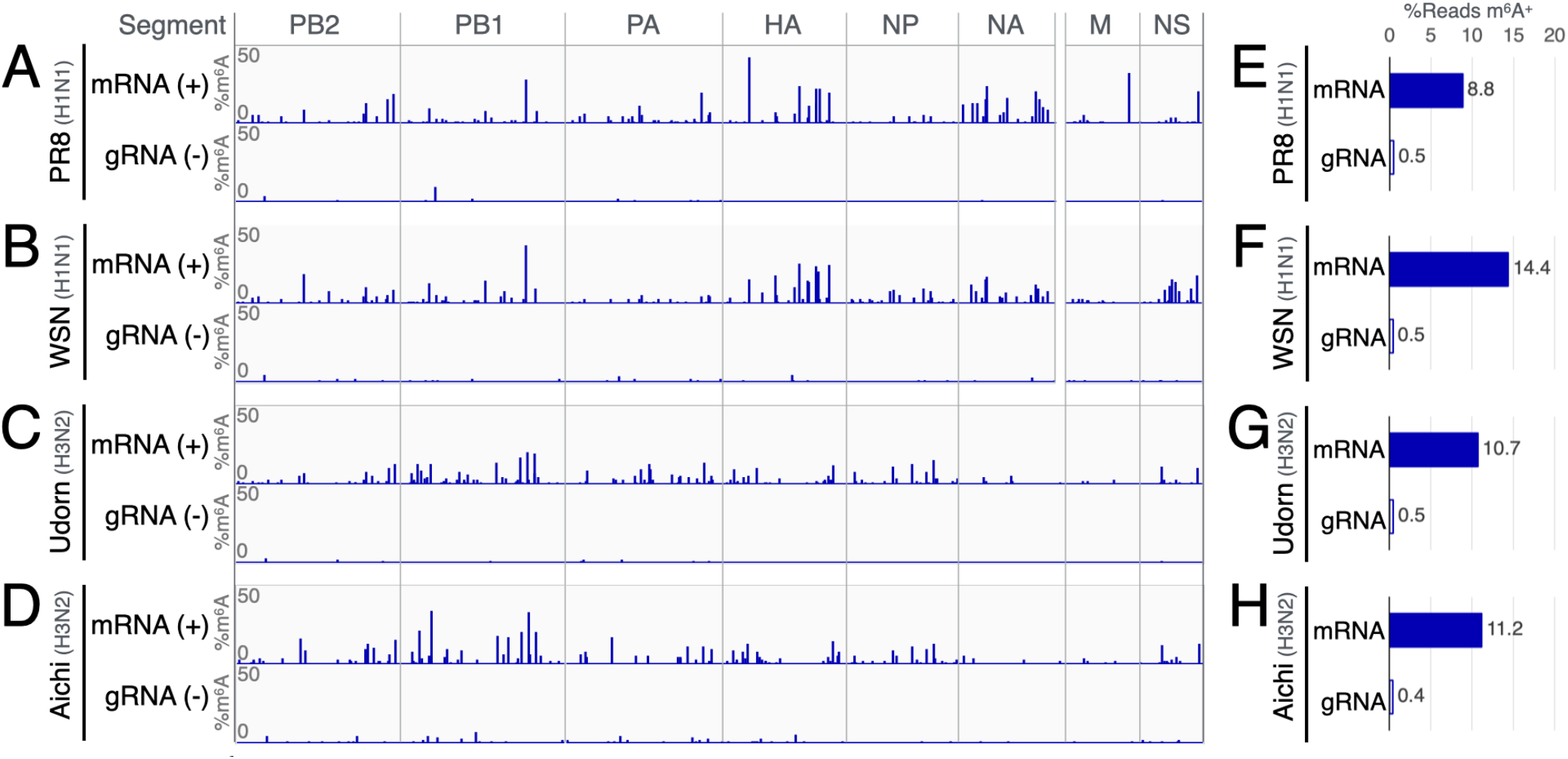
Low m^6^A methylation of viral antisense/genomic RNA is conserved across multiple strains. **(A-D)** The stoichiometry and location of m^6^A methylations mapped via Nanopore DRS across two H1N1 (PR8 & WSN) and two H3N2 strains (Udorn & Aichi), with the % methylation (%m^6^A) on the sense (mRNA (+)) and antisense (gRNA (-)) strand of each viral genome shown in blue vertical bars. **(E-H)** Percent of viral sense (mRNA) and antisense (gRNA) reads with at least one m^6^A detected, shown separately for each viral strain.

### Artificial addition of m^6^A on IAV genomic RNAs enhance NP binding slightly yet does not enhance viral production in cells

A recent study mapped the m^6^A sites on an avian influenza H5N6 isolate A/Goose/Sichuan/SC15/2015 (SC15) using meRIP-seq (15), and reported that viral nucleoprotein (NP) binds genomic RNA segments inefficiently when the meRIP-mapped m^6^A sites were mutated. If m^6^A on genomic RNA were indeed required for packaging of these RNA into vRNPs, we would expect an enrichment of m^6^A on the virion packaged genomic RNAs. Yet, Figs. 1-2 show very little m^6^A on the antisense genomic RNA across four IAV virus strains, regardless if we assayed the intracellular antisense genomic RNA or virion-extracted RNA. In light of this low m^6^A occurrence, we proceeded to test if artificially increasing the total amount of m^6^A may enhance vRNP packaging and viral replication. To this end, we cloned the HA and NS segments of PR8 behind a T7 phage promoter and *in vitro* transcribed the genomic RNA of these two segments in the antisense direction using either ATP or m^6^ATP in the mix of transcription ingredients, which would produce HA and NS genomic RNA either completely free of any methylation or with all adenosines methylated as m^6^A (Fig. 3A). NP protein were immunoprecipitated from NP-expressing cell lysates, and co-incubated with the artificial m^6^A+ and modification-free genomic RNA segments. Upon quantifying the NP pulled down RNA, we observed a slight ∼25% increase in NP binding of fully m^6^A+ genomic RNA over unmethylated RNA (Fig. 3B). However, considering that naturally produced RNA is almost never 100% m^6^A+, we alternatively tested the impact of m^6^A using an A549 cell line that carries an exogenous inducible copy of the m^6^A methyltransferase METTL3, which upon doxycycline treatment will overexpress METTL3 (Fig. 3C), and increase m^6^A methylation (Fig. 3D). Infection of these “m^6^A-high” A549 cells with IAV, does not lead to any notable increase of viral genomic RNA or mRNA at 16hrs post infection (Fig. 3E-F), nor does it lead to any significant change in viral production titers as determined by plaque assay (Figs. 3G-H). Thus, while artificially produced fully m^6^A methylated IAV genomic RNA may slightly enhance NP binding, it is unlikely that the low occurrence of m^6^A on normal IAV genomic RNA can be boosted *in cellulo* to a methylation rate that may functionally support viral production.

**Fig 3.**
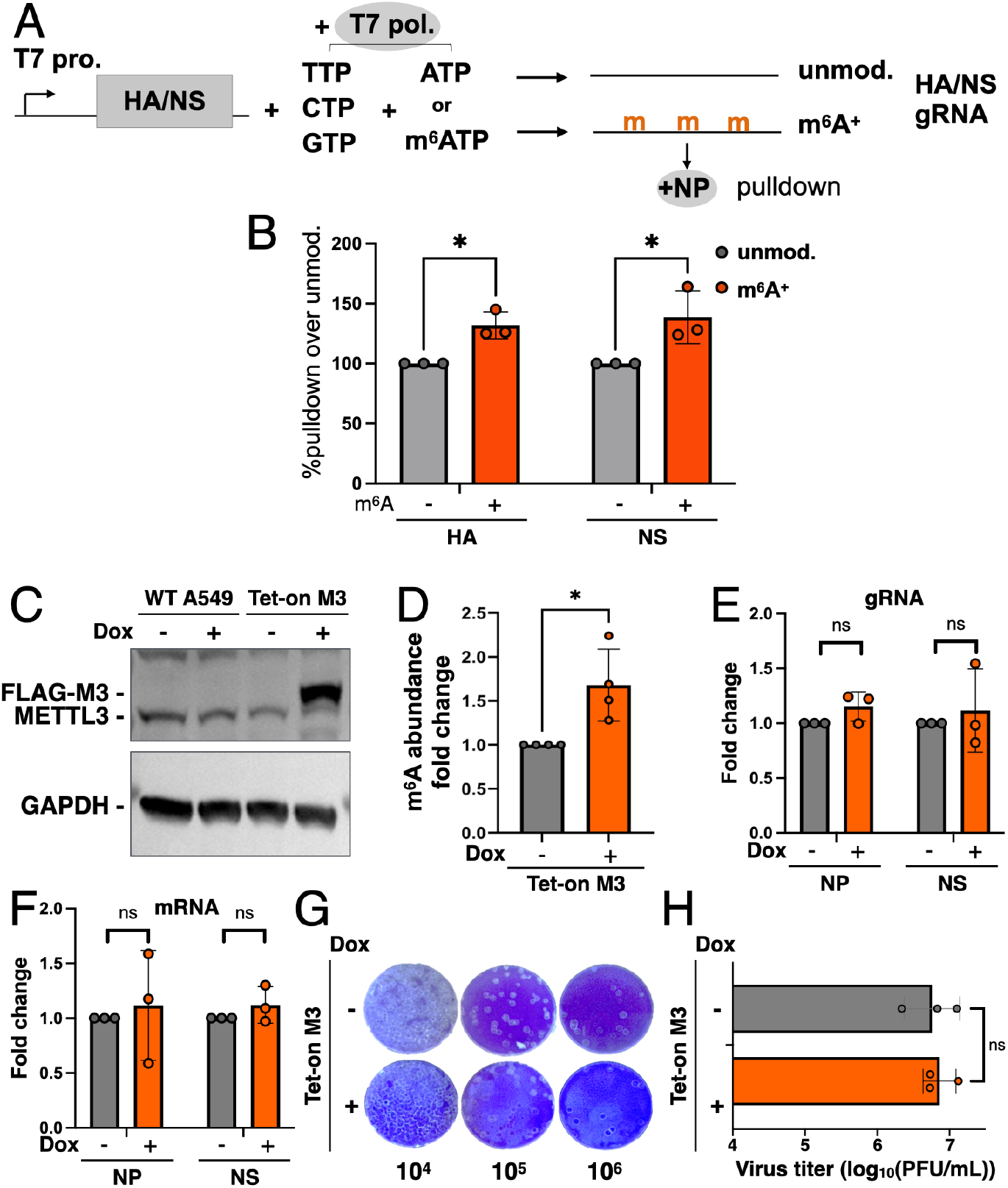
Addition of m^6^A onto viral gRNA does not substantially increase viral packaging. The HA and NS segments of IAV gRNA were *in vitro* transcribed (IVT) with or without the addition of m^6^A and co-incubated with NP protein for RNA pulldown analysis, as summarized in **(A)** and the NP-pulled down RNA quantified by qPCR **(B)**. Doxycycline (Dox) -induced FLAG-tagged METTL3 (M3) overexpression was confirmed by Western blot **(C)** and the resulting m^6^A levels measured by ELISA **(D)**. Cells with or without Dox-induced M3 overexpression were infected with influenza A virus (IAV, PR8) at a multiplicity of infection (MOI) of 1 for 16hrs, and the resulting levels of IAV genomic RNA (gRNA) and mRNA quantified via qPCR **(E-F)**, with the supernatant viral titers determined by plaque assay **(G-H)**. Statistical analyses for panels E&F done by two-way ANOVA, all others by two-tailed Welch’s T test, error bars=SD, *p<0.05.

### m^6^A supports IAV mRNA levels and overall replication

As we could not confirm the presence nor function of m^6^A located on the antisense genomic RNA, we next seek to confirm if m^6^A is indeed proviral for IAV as previously reported. To this end, we transiently knocked down the m^6^A methyltransferase complex enzymatic subunit METTL3 via RNA interference (RNAi), which depleted the METTL3 protein levels by more than 90% (Fig. 4A & 4C), and diminished all tested viral proteins and mRNAs to almost undetectable levels after 48hrs of spreading infection (Figs. 4A-B). In a single round infection test, we also observed a significant >50% loss of viral proteins and mRNAs 24hrs post infection (Figs. 4C-D), suggesting that the loss of METTL3 negatively impacted viral replication not by limiting spread but mostly by diminishing the production of viral mRNA. A plaque assay of virus-containing supernatants from infected control (siCtrl) and METTL3 knockdown cells (siM3) showed almost 10-fold less viral production from siM3 cells than that from controls (Fig. 4E), confirming that METTL3 indeed is required for maintaining efficient viral replication. If METTL3 and its’ m^6^A-deposition activity is indeed required for viral replication, we would expect this to be a viable antiviral strategy. To this end we tested, as a proof of concept, the small molecule inhibitor STM2457, which binds and inhibits the methyltransfer enzymatic core of METTL3 with high specificity (34). As predicted, IAV infection of A549 cells treated with non-lethal dosages of STM2457 (Fig. 4G, gray dashed line), indeed lead to a dosage dependent loss of viral protein and mRNA levels (Figs. 4F-G). We thus conclude that METTL3-dependent m^6^A methylation is indeed proviral as previously reported, and that this can be a viable antiviral target.

**Fig 4.**
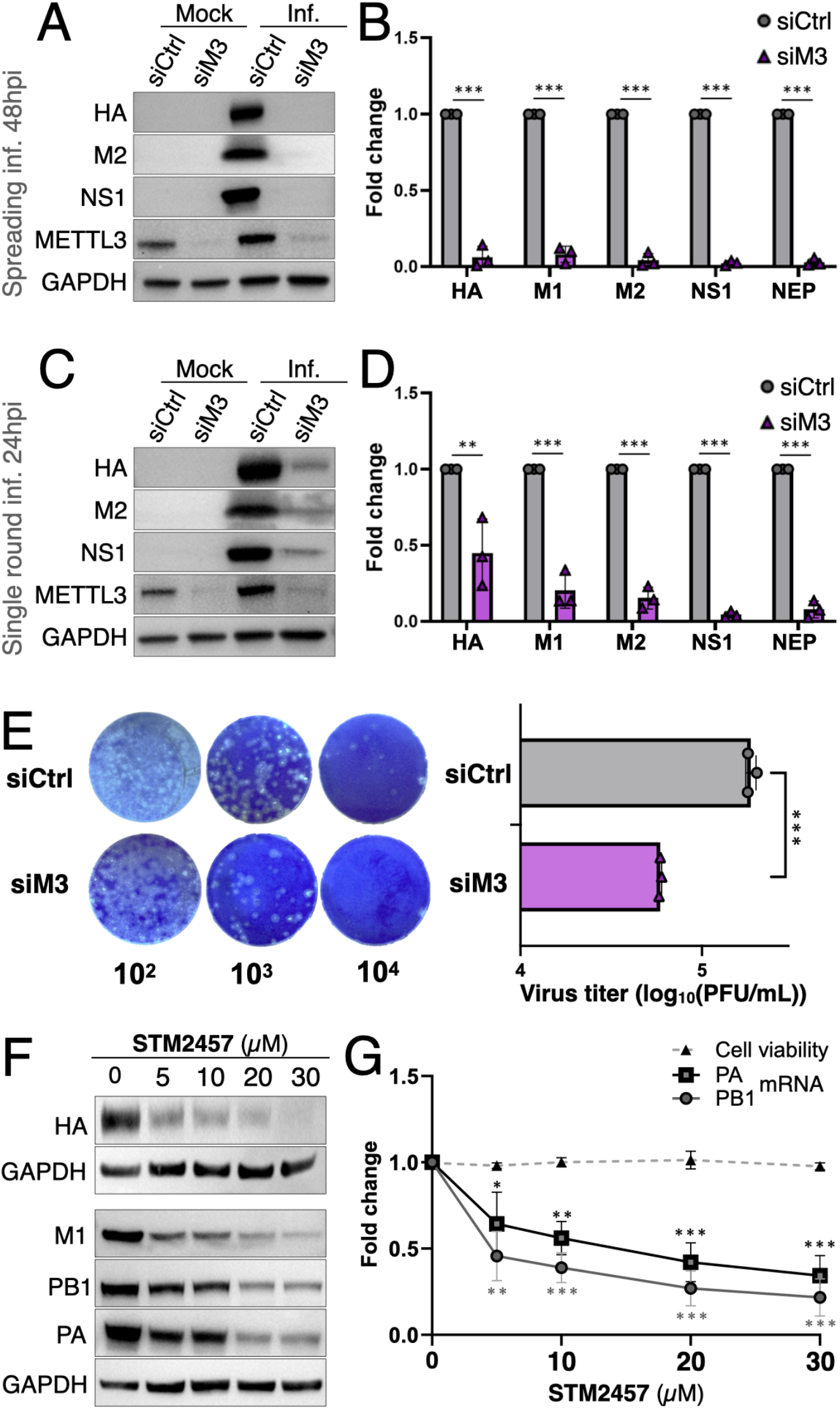
m^6^A ensures efficient gene expression from IAV mRNA. siMETTL3 (siM3) or control (siCtrl) -treated cells were infected with IAV (PR8) in a spreading infection (MOI = 0.1) and analyzed at 48hrs post infection (hpi) for viral protein production by Western blot **(A)** and RNA production quantified via qPCR **(B)**. siM3 and siCtrl-treated cells subject to a single round infection (MOI = 0.5) were also collected 24hpi, and analyzed by Western blot **(C)** and qPCR **(D)**. Viral production titers with or without METTL3 knockdown were determined by plaque assay **(E)**. A549 cells treated with various levels of METTL3-inhibitor STM2457 were infected (PR8) at an MOI of 0.1 and collected 24hpi, and assayed for viral protein **(F)** and mRNA production levels **(G)**. Cell viability under drug treatment also shown as dashed line **(G)**. Statistical analyses for panels B&D done by two-way ANOVA, all others by two-tailed Welch’s T test, error bars=SD, *p<0.05, **p<0.01, ***p<0.001.

### m^6^A enhancement of viral mRNA is dependent on the m^6^A reader YTHDF2 enhancing viral RNA stability

The replication of IAV was previously reported to be enhanced by m^6^A via its’ recognition by the m^6^A reader protein YTHDF2, with the mechanism unknown. With minimal m^6^A present on the genomic RNA making it unlikely to enhance genomic RNA packaging, we sought to elucidate the mechanism behind how m^6^A and YTHDF2 supports viral mRNA levels. We first utilized CRISPR-Cas9 to knockout YTHDF2 from A549 cells (ΔY2), which indeed led to a significant loss of viral protein and mRNA levels (Fig. S2A-B). However, our CRISPR-edited ΔY2 A549 cells consistently lose this phenotype within two weeks of single cell cloning, suggesting the likely development of compensatory mutations. We thus had to switch to an RNAi-mediated transient knockdown approach. Transient knockdown of YTHDF2 using small interfering RNAs (siRNAs) indeed led to a significant loss of viral protein and mRNA expression across spreading (Figs. 5A-B) and single round infection conditions (Figs. 5C-D), suggesting that m^6^A and YTHDF2 regulates viral replication on the mRNA level. m^6^A generally induces the degradation of host mRNAs, while our previous findings suggest that m^6^A can instead prevent the premature degradation of HIV-1 viral RNA (3, 8, 35). We thus proceeded to test if m^6^A may also stabilize IAV RNA. We first treated infected cells with Actinomycin D (ActD) to block RNA production. ActD mainly blocks the host DNA dependent RNA polymerase (RNA polymerase II), while IAV mRNA production is instead dependent on the viral RNA polymerase. We reasoned that since viral RNA polymerase snatches 5’ caps from RNA polymerase II -generated host mRNAs, ActD can indirectly block viral mRNA production, albeit with some delay. Upon quantifying the residual viral mRNA levels of two representative transcripts, we observed these viral mRNAs decaying significantly faster in cells with YTHDF2 knocked down (Figs. S2C-D). As ActD blocks host mRNA production as well, we next utilized the IAV RNA polymerase inhibitor Baloxavir which specifically blocks viral mRNA production while minimizing effects on the host cell (36). Upon treatment of infected cells with Baloxavir, and quantifying the residual amounts of viral mRNA over time, we confirm that knockdown of YTHDF2 led to a statistically significant faster decay of viral mRNAs (Figs. 5E-F). Considering that YTHDF2 may regulate the decay of m^6^A^+^ RNA by interacting with poly(A) tail deadenylase complexes (37), and that removal of the poly(A) tail is the rate determining step for the decay of most mRNAs (38), we additionally measured the poly(A) tail length of IAV mRNAs via Nanopore direct RNA sequencing. Unexpectedly, we observed a larger proportion of viral RNA carrying poly(A) tails >90nt in YTHDF2 knockdown cells than in control cells (Fig. 5G). As long tails should better protect RNA from 3’end-initiated degradation yet YTHDF2 knockdown led to decreased viral RNA levels and faster RNA decay (Figs. 5D-F), YTHDF2-mediated viral RNA stabilization likely works through a poly(A) tail -independent mechanism.

**Fig 5.**
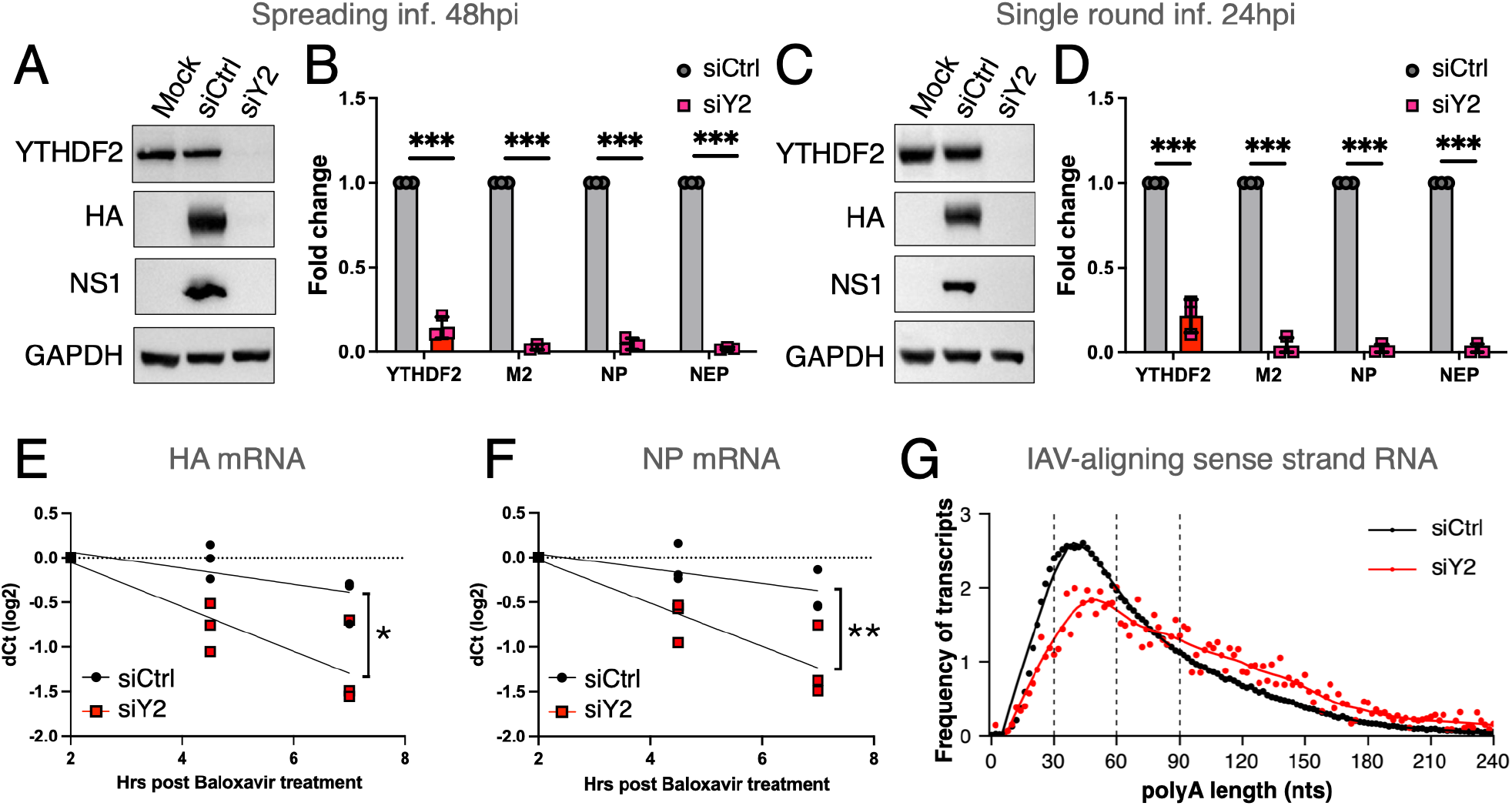
The m^6^A reader YTHDF2 ensures the stability of IAV mRNA through a 3’ terminal -independent mechanism. siYTHDF2 (siY2) or control (siCtrl) -treated cells were infected with PR8 (MOI = 0.1), subjected to a spreading infection and analyzed at 48hpi for viral protein production by Western blot **(A)** and RNA production quantified via qPCR **(B)**. siY2 and siCtrl-treated cells subject to a single round infection (MOI = 1) were also collected 24hpi, and analyzed by Western blot **(C)** and qPCR **(D)**. Similarly, siY2 or siCtrl cells under 1 MOI single round infections were subjected to IAV-specific transcription inhibition with Baloxavir and collected at the indicated time points for qPCR analysis of viral HA and NP mRNA decay **(E-F)**. Poly(A) tail length analysis of viral mRNAs from siCtrl and siY2 -treated cells as determined by Nanopore direct RNA sequencing **(G)**. Statistical analyses done by two-way ANOVA, error bars=SD; slopes of RNA decay regression lines compared by analysis of covariance (ANCOVA); *p<0.05, **p<0.01, ***p<0.001.

## Discussion

Given the profound impact of Influenza A virus on human health, the molecular biology of this virus has been extensively studied. It was thus not surprising that this virus happens to be one of the first few viruses whose RNA was probed for the presence of the non-cap RNA methylation, m^6^A. Past and present reports have added up to a consensus that m^6^A is present on IAV RNA, and that this modification can functionally enhance viral replication through multiple mechanisms, including increased gene expression, fine tuning of the splice rate of the NS genes, and facilitating the packaging of the antisense genomic RNA (12–15, 39). However, studies of each era also reflected the limitations of contemporary techniques of the time, with studies in the 1970s leaving the function and location of m^6^A unclear, while studies in the past decade were all based on immunoprecipitations using m^6^A-targeting antibodies which obscures the stoichiometry of each m^6^A site. Here, exploiting the latest m^6^A mapping techniques, we propose a series of updates to the current understanding of how IAV is regulated via m^6^A. First of all, m^6^A is mostly present on the sense-strand mRNAs, while m^6^A on the antisense genomic RNA was only detected at low abundances that are unlikely to have an impact on vRNP packaging or viral replication in general (Figs. 1-3 & S1). In light of this observation, we confirm that m^6^A and the associated reader protein YTHDF2 are proviral through the enhancement of viral mRNA stability, and that this is most likely explained by the m^6^A located on viral mRNAs (Figs. 4-5).

While probing the modification landscape of IAV viral RNAs, we initially compared that of infected A549 mRNA (including host and viral mRNAs) and RNA purified from embryonated chicken eggs -produced virions, using UPLC-MS/MS, and observed a much lower %m^6^A/A rate in virion RNA versus A549 intracellular mRNAs (Figs. 1A-B), leading us to hypothesize a difference in the propensity of avian and human m^6^A methyltransferases to methylate IAV RNA. However, further probing of the m^6^A on A549-produced virion RNA and the intracellular mRNA of infected chicken cells ruled out this hypothesis and instead suggested that the genomic RNA are sparsely methylated and that previous observations of genomic RNA methylation were likely due to PCR amplification of the rare population of methylated RNA during sequencing library preparation for meRIP. Follow up detection of the m^6^A on genomic RNA using mapping techniques that more accurately reflect the stoichiometry such as Nanopore direct RNA-seq and GLORI-seq indeed supports this view (Figs. 1C-F & 2), suggesting that m^6^A sites on the genomic RNA are few and far in between, with most sites 1-2% methylated and no sites >5%.

A recent report has proposed that m^6^A on the genomic RNA on the avian flu isolate SC15 is required for coating the viral genomic RNA with the viral nucleoprotein (NP) and packaging into vRNPs (15), a mechanism that would predict the virion packaged RNA to be enriched with m^6^A. However, our data suggests that the virion RNA of viral isolate PR8, as well as the intracellular genomic RNA of WSN, Udorn and Aichi are all sparsely methylated (Figs. 1-2). While we do not have access to the SC15 isolate and thus cannot rule out differences between isolates, our data suggest that m^6^A is unlikely to be universally important for the packaging of all influenza isolates. While SC15 is an avian isolate, growing PR8 in chicken cell and eggs do not result in increased methylation of viral genomic RNA (Fig. 1A-B), making it unlikely to be an avian host -specific phenomenon. Lastly, manually increasing the methylation on PR8 genomic RNA resulted in a minor 25% boost of *in vitro* NP binding when we raised the m^6^A methylation rate to 100% (Fig. 3). Considering that the average cellular mRNA m^6^A/A rate is in the ∼0.2-0.6% range, it is very unlikely that any host can provide an m^6^A-dependent boost substantial enough to enhance vRNP production. Thus, our data does not support m^6^A as an essential factor for viral vRNP packaging. Notably, data supporting the need of m^6^A for NP binding were mostly generated through point mutations of meRIP-mapped m^6^A sites (15). If each of these mutated m^6^A sites were initially methylated on <5% of viral transcripts, this would mean the mutations would be introducing methylation-unrelated changes to the other 95% of viral genomic RNA molecules, the impacts of which hard to interpret unless the function of those specific altered nucleotides on the SC15 vRNPs were further characterized.

In contrast to that on genomic RNAs, our data suggests that the majority of viral m^6^A is on mRNA. Multiple m^6^A site are consistently found via PA-m^6^A-seq and Nanopore direct RNA-seq, with those on PR8 also confirmed by GLORI (Figs. 1-2 & S1). We note that GLORI detected less m^6^A sites than Nanopore as the standard GLORI analysis pipeline removes sites <10% m^6^A positive to avoid false positives (28), nonetheless all the major sites on IAV mRNA were confirmed. In light of this, the proviral function of m^6^A is most likely due to the methylations on the mRNA instead of the genomic RNA, which is also supported by the fact that METTL3 and YTHDF2 knockdowns strongly impact viral RNA and protein production even in single round infections where viral spread was not assayed (Figs. 4C-D & 5C-D). To this end, our data suggest that m^6^A may protect viral RNAs from premature degradation (Fig. 5), in a likely evolutionary convergence with our previous observation on HIV-1 m^6^A. The presence of m^6^A induces the degradation of the vast majority of host mRNAs with some exceptions, yet we found m^6^A to instead prevent degradation of HIV-1 RNA (3, 8, 35, 40). Our observation here that m^6^A also protects IAV mRNAs from degradation, while contrary to the reported function of m^6^A on cellular RNA, does make sense considering that the rapid evolution of viruses such as IAV and HIV-1 should only retain m^6^A if it is evolutionarily beneficial (5). It would also be interesting to speculate the potential presence of a hypothetical RNA-targeting antiviral restriction factor that m^6^A may protect against. In search for clues of this restriction factor, we considered that m^6^A has been reported to regulate the poly(A) tails of mRNAs and the shortening of poly(A) tails is generally rate limiting for RNA decay (37). We thus utilized Nanopore direct RNA-seq to elucidate the poly(A) tail lengths of IAV mRNA (Fig. 5G). However, we see the opposite of what we would expect if YTHDF2/m^6^A protection of viral mRNA stability involves protecting poly(A) tail length, instead showing longer viral mRNA tails when YTHDF2 is knocked down. Thus, it is likely that m^6^A protects viral mRNAs from a host restriction mechanism that is not related to poly(A) tails and may instead attack viral RNAs from the 5’end or internal regions.

As m^6^A is universally proviral for a wide variety of viruses, we have previously speculated that m^6^A-targeting inhibitors may serve as broad-spectrum antivirals (5). Indeed, the m^6^A methyltransferase METTL3 inhibitor STM2457 was previously shown to inhibit the replication of the human coronaviruses OC43 and SARS-CoV-2, as well as hepatitis B virus (HBV) (7, 41). Here, we demonstrate that STM2457 can similarly be antiviral against IAV (Figs. 4F-G), broadening the antiviral potential of this inhibitor. As we work towards elucidating antiviral restriction mechanisms that are blocked by the presence of m^6^A on viral RNAs, we hope to unveil new antiviral targets that may be more specific to viral RNAs and minimally impact host RNA.

## Data availability

All deep sequencing data have been deposited at the NCBI GEO database under accession number GSE344571.

## Acknowledgements

We thank Peter Palese for kindly providing us with reverse genetics plasmids for the generation of PR8 virus stocks, Alexander Price and Matthew Weitzman for generously providing us with METTL3 knockout and tet-on inducible METTL3 overexpressing A549 cells, as well as Hui-Wen (Winni) Chen for kindly providing us with the DF1 cell line. Special thanks to Storm Therapeutics Ltd. for kindly sharing us with an initial batch of STM2457. We also thank Chengqi Yi and Hanxiao Sun for sharing detailed protocols of GLORI-seq. This work cannot be done without the following core facilities at Academia Sinica: IBMS DNA Sequencing (AS-CFII-111-211), IMB bioinformatics core, IMB genomics core, the Biodiversity Research Center (BRCAS) High Throughput Genomics Core, Academia Sinica grid-computing center (ASGC), and the BioTReC Infectious Disease Core (AS-NBRPCF-114-201_55FC). This research was supported by the NSTC grant 111-2320-B-001-006-MY2 to K.T., Academia Sinica career development award AS-CDA-112-L02 to K.T. and the Intramural Research Program of the NIH, National Institute of Environmental Health Sciences (project ZIA ES103355) to M.M. The contributions of the NIH author(s) are considered Works of the United States Government. The findings and conclusions presented in this paper are those of the author(s) and do not necessarily reflect the views of the NIH or the U.S. Department of Health and Human Services.

## Supplemental Material

**Fig. S1.**
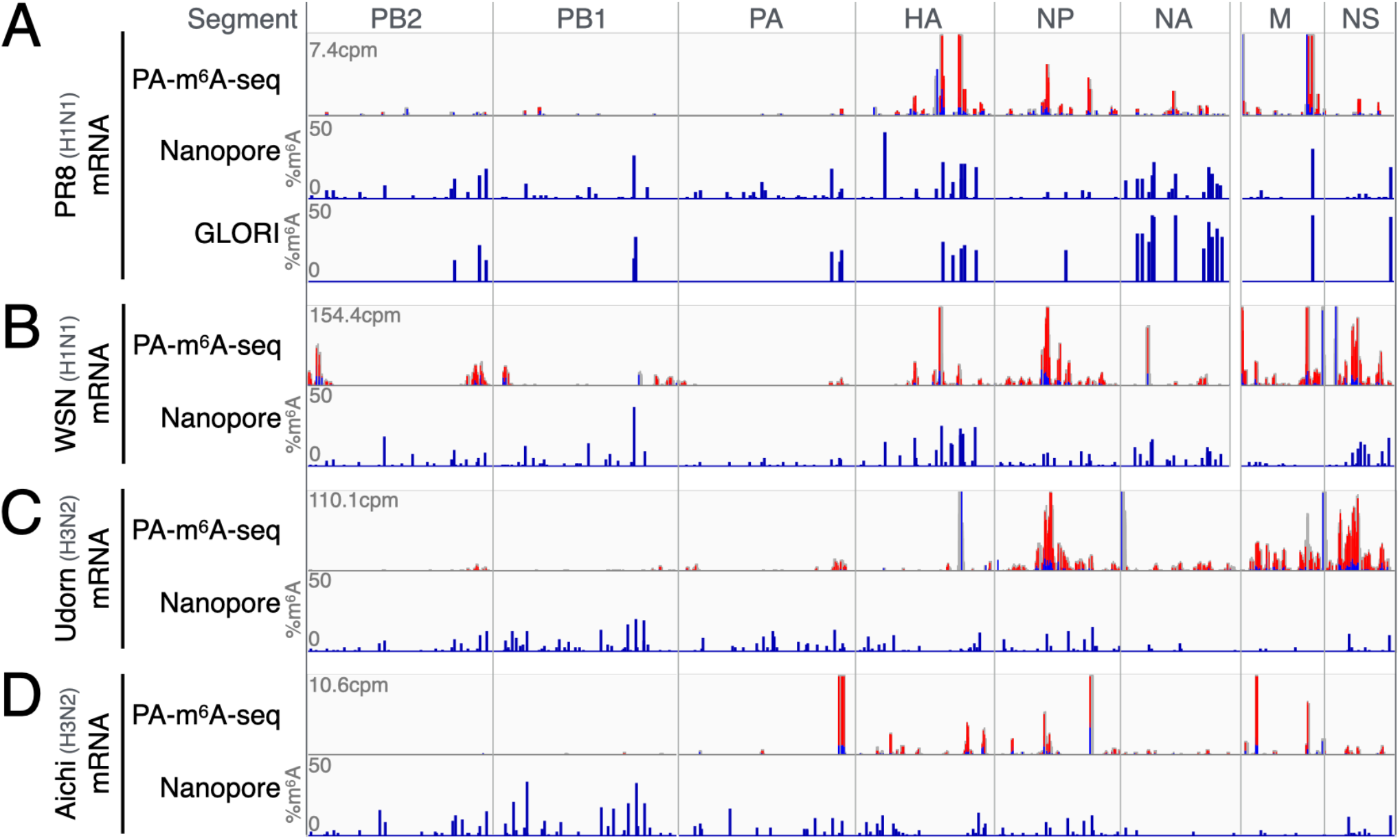
m^6^A sites on IAV mRNA consistently detected by multiple methods. The Nanopore-detected m^6^A sites of four IAV strains as shown in Fig. 2, compared with m^6^A antibody-immunoprecipitated sites (PA-m^6^A-seq), as well as the GLORI-mapped sites as shown in Fig. 1E. Red & blue bars in the PA-m^6^A-seq lanes indicate sites of antibody-RNA crosslinking-induced T>C mutations, characteristic of 4SU-mediated photo activated crosslink and immunoprecipitation (CLIP) products.

**Fig. S2.**
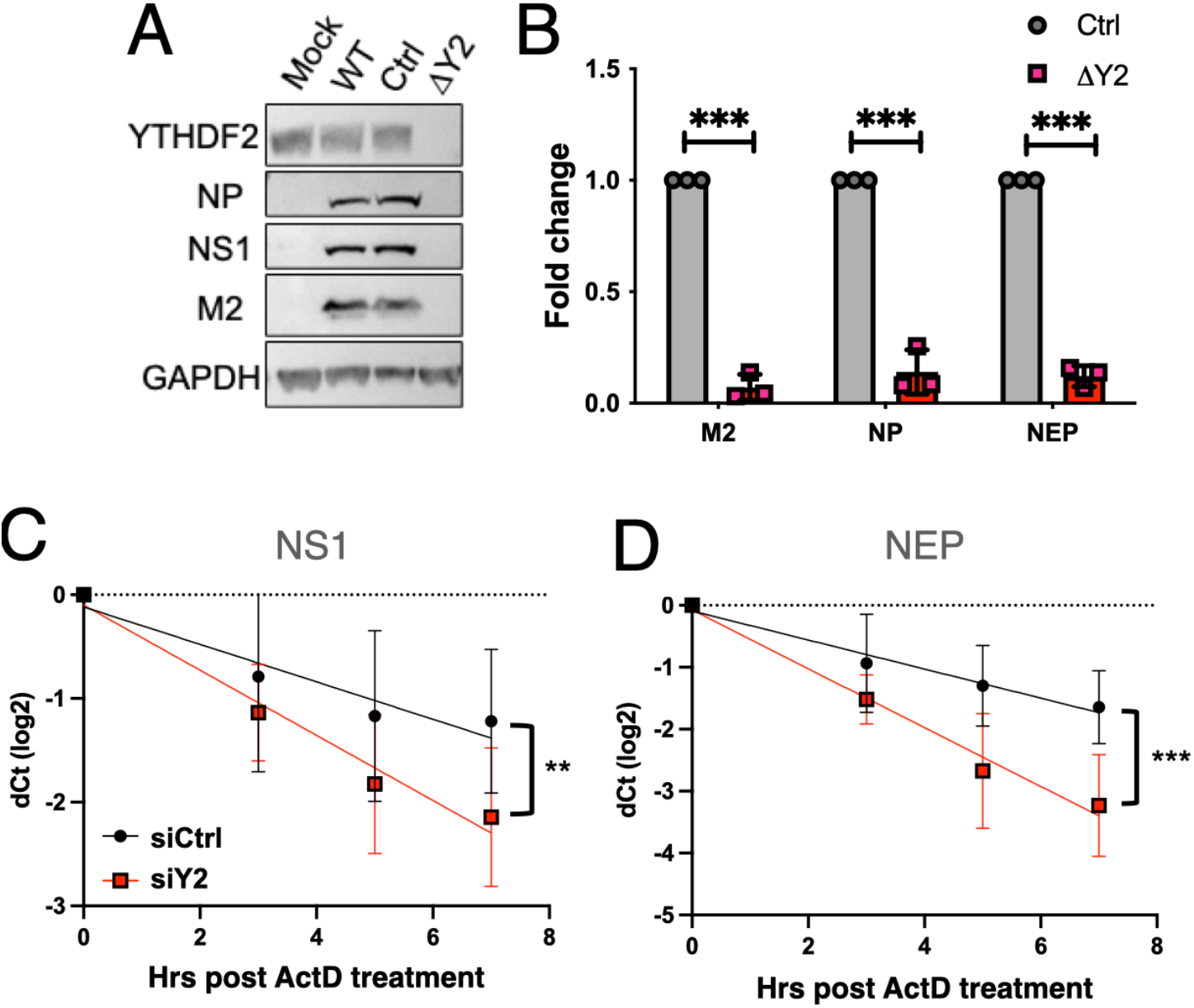
The m^6^A reader protein YTHDF2 stabilizes IAV mRNAs. A549 cells, either untransduced (WT), transduced with Cas9 with a non-targeting guide RNA (Ctrl) or with Cas9 and a YTHDF2-targeting guide RNA (ΔY2), were infected with PR8 (MOI = 0.1), subjected to a spreading infection and analyzed at 48hpi for viral protein production by Western blot **(A)** and RNA production quantified via qPCR **(B)**. Similar to Figs. 5E-F, siY2 or siCtrl -transfected A549 cells under 1 MOI single round infections were subjected to general transcription inhibition using Actinomycin D (ActD), and the residual viral RNA quantified by qRT-PCR at various time points after transcription block **(C-D)**.

